# CRISPR/Cas9-mediated transformation enables functional characterization of the effector *Avr4* in the banana pathogen *Pseudocercospora fijiensis*

**DOI:** 10.64898/2026.08.10.742443

**Authors:** Maikel B.F. Steentjes, Gregory Ashe, Patricia Schöppl, Rahim Mehrabi, Gert H.J. Kema

**Author notes:** Laboratory of Biochemistry, Wageningen University & Research, Stippeneng 4, 6708 WE, Wageningen, the Netherlands. Keygene N.V., Wageningen 6700 AE, The Netherlands.

## Abstract

*Pseudocercospora fijiensis* is the causal agent of Black Leaf Streak Disease (BLSD), also known as black Sigatoka, in banana. The disease affects many banana varieties, including the highly susceptible Cavendish banana that dominates global production and the export trade, and several cooking bananas that are a staple food for hundreds of millions of people worldwide. Currently, the disease is controlled using preventative fungicide treatments with up to 70 applications per year in Cavendish plantations, which accounts for approximately 30% of the production costs. Resistant cultivars are required for more sustainable production, but no resistance gene to BLSD has been identified. This is partly due to the poor genetic amenability of *P. fijiensis* and the lack of methods for functional gene analysis.

To address these limitations, we developed a CRISPR/Cas9-mediated transformation system specifically optimized for *P. fijiensis*. We established a protocol to produce protoplasts, evaluated their capacity to regenerate into new colonies, and assessed antibiotic sensitivity. Subsequently, we confirmed the integration of foreign DNA, including resistance markers, using PEG-mediated transformation. We demonstrated targeted transformation using CRISPR-Cas9 to knockout the polyketide synthase gene *PKS10-1,* which is responsible for the production of the pigment melanin, and the mitogen-activated protein kinase (MAPK) gene *Fus3*.

Following the successful generation of knockout mutants for these genes, achieving gene targeting efficiencies of respectively 96% and 58%, we subsequently generated knockout mutants of the renowned effector *Avr4* in *P. fijiensis*. The resulting mutants exhibited no reduction in virulence on the susceptible banana cultivar Cavendish. In addition, we used the wild-type isolate and *Avr4* knockout strains to test the resistant banana accession Calcutta 4. Contrary to a previous study, we demonstrate that *Avr4* does not explain the resistance of Calcutta 4, suggesting that resistance is instead triggered by the recognition of other hitherto unknown effectors. The established CRISPR/Cas9-mediated disruption system is highly efficient and enables routine functional gene characterization, which will help to elucidate genes involved in banana-*P. fijiensis* interaction, thereby supporting the discovery of resistance genes against BLSD.

## Introduction

Banana is a diverse crop that comprises the popular dessert fruit, as well as several types of cooking bananas. These cooking bananas are the fourth most important staple crop following rice, wheat, and maize, and they are cultivated in over 130 countries primarily located in tropical and subtropical climates (Arias, 2003; Churchill, 2011; Evans et al., 2020). Despite the vast genetic diversity across banana germplasm, historically, banana cropping for international markets has been intimately linked with monocultures. The iconic Gros Michel banana was cultivated throughout Latin America but succumbed over a period of several decades to Fusarium wilt until the epidemic reached a tipping point in the 1950s (Ploetz, 2005; Ploetz et al., 2015). Henceforward, Cavendish varieties gradually replaced collapsed Gros Michel plantations due to their excellent resistance to Fusarium wilt. Since then, Cavendish varieties have been planted in many more countries and now dominate the global production (>50%) and the export trade (>95%) (García-Bastidas et al., 2022). Evidently, these vast monocultures make contemporary banana production - again-extremely vulnerable to diseases (Ploetz et al., 2015; Drenth and Kema, 2021). As a result, banana production is currently heavily affected by several pests and diseases, such as Black Leaf Streak Disease (BLSD), or black Sigatoka (Drenth and Kema, 2021; Drenth and Kema, 2024).

The disease is caused by the ascomycete fungus *Pseudocercospora fijiensis* and is arguably the costliest and most destructive foliar disease of banana cultivation worldwide (Marin et al., 2003; Alakonya et al., 2018; Noar et al., 2022). It affects almost all banana varieties, including the Cavendish banana and most cooking-banana cultivars (Pasberg-Gauhl and Gauhl, 1996; Marin et al., 2003; Churchill, 2011; Drenth and Kema, 2024), and causes severe foliar necrosis which reduces photosynthesis, leading to major losses in banana yield and fruit quality, including smaller, lighter bunches and poorly filled fruits. The fruits of infected plants also show premature and uneven ripening, making them unsuitable for shipment and export. Yield losses can exceed 38% in plantains and over 50% in export bananas when unmanaged (Marin et al., 2003; Noar et al., 2022). Currently, the main control measure to manage BLSD is frequent fungicide applications. To effectively treat the vast monocultures, fungicides are sprayed by aircraft up to 70 times per year (Chong et al., 2011; Ploetz et al., 2015; Chong et al., 2024). This management strategy is extremely unsustainable due to the high economic and environmental burden. Therefore, the need for resistant banana cultivars is higher than ever. However, banana breeding is complex. This is partly due to the genetically diverse origin of the plant species, its seedlessness and sterility (Perrier et al., 2011; Alakonya et al., 2018; Batte et al., 2019; Smith et al., 2020). On the other hand, since the fruit developed over the last century from an exquisite luxury into a commodity, many stakeholders are also involved in the logistics, ripening, and retail, which complicates the introduction of new cultivars. Nevertheless, presently, BLSD is - next to Fusarium wilt-considered to be the major threat of the entire industry (Drenth and Kema, 2024) and urgently requires sustainable solutions for disease control.

In the search for alternative strategies to combat BLSD, fundamental knowledge about the biology and pathogenicity of *P. fijiensis* is essential. Progress in understanding the molecular mechanisms underlying pathogenicity has been slow, since *P. fijiensis* is recalcitrant due to its poor experimental amenability (Arango Isaza et al., 2016; Alakonya et al., 2018; Diaz-Trujillo et al., 2018). Mycelial colonies melanize quickly, and it is virtually impossible to routinely produce high numbers of conidia. Therefore, disease assays rely on mycelial fragments and usually take 5-6 weeks. Consequently, the development of efficient genetic transformation methods that are required for deciphering the pathogenicity of *P. fijiensis* has proven challenging. Several methods have been explored, including Polyethylene Glycol (PEG)-mediated transformation (Balint-Kurti et al., 2001; Portal et al., 2012) and *Agrobacterium tumefaciens*-mediated transformation (Onyilo et al., 2017; Diaz-Trujillo et al., 2018) but both transformation and gene targeting efficiencies remained very low. Alternative strategies such as transformation based on underwater shock waves, resulted in better transformation efficiencies (Escobar-Tovar et al., 2015), but taken together, all approaches suffer from a lack of precision: an unpredictable number of largely off-target insertions. Hence, these methods are unsuitable for reliable gene knockouts in the quest for the identification and functional characterization of virulence factors.

The emergence of clustered regularly interspaced short palindromic repeats (CRISPR)-associated RNA-guided Cas9 endonuclease technology has revolutionized genetic research (Wright et al., 2016). CRISPR originated as an adaptive immune mechanism in bacteria, protecting against invading viruses. In this system, the Cas9 endonuclease forms a complex with a single guide RNA (sgRNA) that directs sequence-specific recognition and introduces a double-strand break at the target DNA locus (Jinek et al., 2012; Cong et al., 2013). Because the sgRNA can be easily redesigned, CRISPR/Cas9 enables precise cleavage at defined genomic sites (Mali et al., 2013). Following DNA cleavage, DNA repair machinery is activated to restore genome integrity. Repair occurs either through non-homologous end joining (NHEJ), which often introduces small insertions or deletions, or through homology-directed repair (HDR) when a repair template is provided (Sander and Joung, 2014; Maruyama et al., 2015). The latter pathway can be exploited to achieve precise integration of foreign DNA sequences into the host genome at the cleavage site. As a result, CRISPR/Cas9 has become a powerful gene-editing tool that is both highly precise and versatile. Importantly, CRISPR/Cas9 has been shown to overcome the low gene-editing efficiencies commonly observed in many filamentous fungi using conventional transformation methods (Song et al., 2019). Despite these advantages, achieving robust Cas9 expression and efficient nuclear targeting of both Cas9 and sgRNA via plasmid-based systems remained challenging in many fungal species (Schuster and Kahmann, 2019). However, the direct delivery of Cas9-sgRNA ribonucleoprotein (RNP) complexes has proven highly effective in a range of filamentous fungi, including *Aspergillus fumigatus*, *Magnaporthe oryzae*, *Fusarium oxysporum*, and *Botrytis cinerea*, resulting in improved editing efficiencies and reduced off-target effects (Al Abdallah et al., 2017; Foster et al., 2018; Wang et al., 2018; Leisen et al., 2020).

In this study, we established a highly efficient CRISPR/Cas9-based gene-editing platform for *P. fijiensis* by systematically optimizing each step of the transformation procedure for this experimentally challenging species. Using an RNP-mediated strategy, we achieved targeted gene disruption efficiencies between 58% and 96%, across three genes. In addition to the polyketide synthase gene *PKS10-1*, which is essential for melanin production (Noar and Daub, 2016), and the mitogen-activated protein kinase (MAPK) gene *Fus3*, which plays a conserved role in fungal pathogenicity (Xu, 2000; Onyilo et al., 2018), we targeted the effector gene *Avr4*. This effector was previously reported to trigger a hypersensitive-like response upon infiltration in the resistant banana accession Calcutta 4 (Arango Isaza et al., 2016), making it an attractive candidate for functional characterization. Overall, the developed efficient CRISPR/Cas9-mediated transformation system in *P. fijiensis* provides a solid foundation for exploring its pathogenicity and virulence, which contributes to the identification of resistance genes required for durable disease control.

## Materials and Methods

### Fungal culture conditions

The *P. fijiensis* wild-type strain P78 was cultured on potato dextrose agar (PDA, Oxoid) plates in the dark at 25 °C. For routine propagation, mycelium was scraped from plates and transferred to 2 mL microcentrifuge tubes containing two 2 mm glass beads. Samples were homogenized in sterile water using a TissueLyser II (Qiagen) for 1 minute at 30 Hz. The resulting mycelial suspension was used to inoculate fresh plates.

To compare culture conditions for biomass production, mycelium was harvested from fully grown PDA plates (2 weeks old) by removing agar and homogenizing the mycelium in water using an ULTRA-TURRAX tube drive (IKA) at 6000 rpm for 3 x 20 s. The homogenate was filtered through sterile cheesecloth and adjusted to an OD_600_ of 2.0. Aliquots of 25 mL were used to inoculate 50 mL of water, potato dextrose broth (PDB, Difco), Gamborg B5 medium with vitamins (GB5, Duchefa) or malt extract medium (ME, Oxoid) in 250 mL Erlenmeyer flasks. Cultures were incubated at 25 °C and 150 rpm in the dark. Mycelial growth was monitored microscopically every 12 h for up to 72 h.

### Protoplast generation and optimization

Liquid cultures growing for 40 h were harvested by centrifugation (1000 g, 10 min). Pellets were washed with KCl solution (0.6 M KCl, 100 mM sodium phosphate pH 5.8) and resuspended in fresh KCl solution at 0.2 g fresh weight per mL. To test the protoplasting capacity of the different enzyme mixtures, 1 mL aliquots of mycelium suspension were incubated with different enzyme mixtures (Table 1). Enzyme solutions were prepared in 2 mL KCL solution, mixed for 30 min on a rolling platform, and filter sterilized (0.45 μm) before use. Mycelial suspensions were incubated with enzyme mixtures at 28 °C on a rocky 3D shaker (gentle agitation) for 3 h. Protoplasts were filtered through sterile nylon mesh (20 μm) and quantified microscopically using a hemocytometer.

**Table 1.**
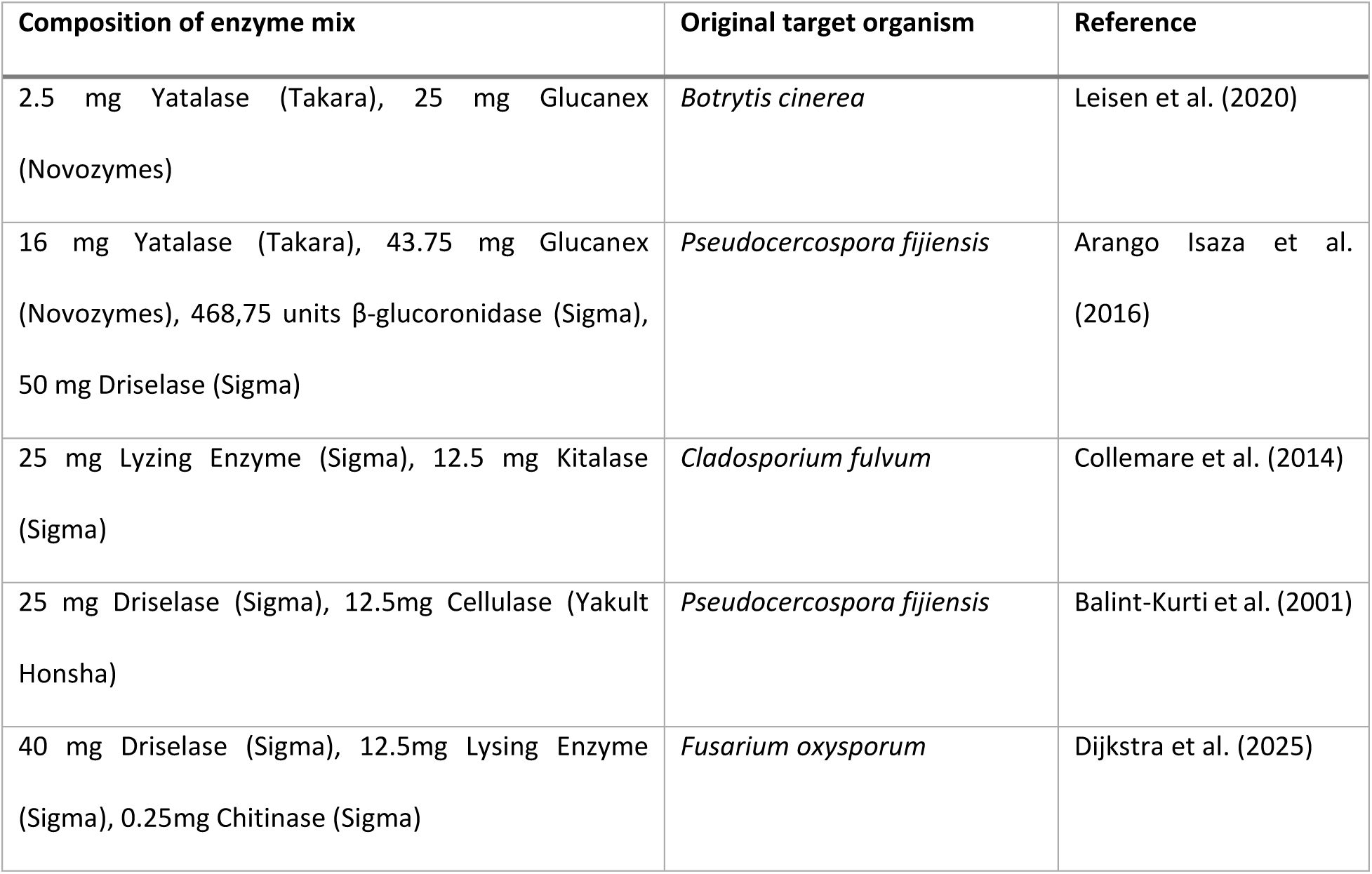
Enzyme mixtures tested for protoplast generation.

To assess the effect of enzyme concentration, total enzyme input was reduced to 50%, 25%, and 12.5% while maintaining constant biomass input. Samples were incubated as described above and protoplast formation was quantified every hour.

To evaluate individual enzyme contributions, each enzyme was tested independently at 20 mg per reaction while maintaining constant biomass input. In addition, omission experiments were performed using the complete enzyme mixture (Arango Isaza et al., 2016), excluding one enzyme at a time.

### Protoplast regeneration and antibiotic sensitivity assays

Freshly generated protoplasts were resuspended in 25 mL molten (42 °C) regeneration medium consisting of Yeast Extract 1 g/L, Casein hydrolysate 1 g/L, Sucrose 275 g/L and agarose 10 g/L, and poured into two petri dishes. After 18 h incubation at 25 °C in the dark, plates were overlaid with 12.5 mL regeneration medium containing increasing concentrations of hygromycin B (0, 12.5, 25, 50, 75 or 100 µg/mL) or nourseothricin (0, 5, 10, 25, 50, 75, 100, 200, 500 or 1000 µg/mL). Plates were subsequently incubated at 25 °C in the dark, and colony formation was assessed after 7 days.

### sgRNA synthesis, RNP assembly, and *in vitro* cleavage assays

sgRNAs were designed and synthesized as described by Leisen et al. (2020). Oligonucleotides used for sgRNA synthesis are listed in Supplemental Table S1. Cas9 nuclease (EnGen Spy Cas9 NLS, New England Biolabs) was used for ribonucleoprotein (RNP) assembly as described by Leisen et al. (2020).

*In vitro* cleavage assays were performed as described by Jinek et al. (2012) with minor modifications. Cas9 (1:1 molar ratio with sgRNAs) was incubated with sgRNAs in 1x r3.1 cleavage buffer (NEB) for 15 min at room temperature. Target DNA was amplified by PCR using the primers listed in Supplemental Table S1, followed by purification. For each reaction, 200 ng target DNA was added to the pre-incubated RNP complex, and reactions were incubated at 37 °C for 1.5 h. Reactions were terminated with 2 mg/mL proteinase K for 20 minutes at 37 °C. DNA fragments were analyzed on a 1% agarose gel in 1x TAE buffer.

### Donor template production

Donor templates were generated by PCR amplification of the hygromycin resistance cassette from plasmid pTel_Hyg (Leisen et al., 2020) using primers containing 60 bp homology arms flanking the target loci (Supplemental Table S1). The size of the PCR products was verified by gel electrophoresis and purified before use in transformation. For *PKS10-1* complementation, the full genomic locus, including the native promoter (951 bp upstream of the start codon) and terminator (1253 bp downstream of the stop codon) was amplified and fused to a nourseothricin resistance cassette amplified from plasmid pTel_Nat (Leisen et al., 2020) using overlap extension PCR. The size of the final product was verified by gel electrophoresis and the product was purified before use in transformation. Fluorescent reporter constructs (GFP and DsRed) were amplified together with the hygromycin resistance cassette from plasmids pCT74:GFP and pCT74:DsRed (Lorang et al., 2001). Also these products were verified by gel electrophoresis and purified before use in transformation.

### CRISPR/Cas9-mediated transformation

Protoplast transformation was adapted from Leisen et al. (2020) with several modifications to optimize the protocol for *P. fijiensis*. Protoplasts were generated as described above and washed two times in ice-cold TMS solution (1 M sorbitol, 10 mM MOPS, pH 6.3) by centrifugation at 1500 g for 5 minutes at 4 °C. Protoplasts were resuspended in cold TMSC buffer (TMS supplemented with 50 mM CaCl_2_) at 5 x 10^5^ – 2 x 10^6^ protoplasts per 100 µl.

For PEG-mediated transformation, 5 µg of donor template DNA per reaction was prepared in Tris-EDTA solution (10 mM Tris-HCl, 1 mM EDTA, 40 mM CaCl2, pH 6.3) and RNP complexes were preassembled. For RNP assembly, 6 μM of EnGen Spy Cas9 NLS enzyme (NEB) and 2 µg sgRNA per target gene were mixed in 1x cleavage buffer r3.1 (NEB) per for each transformation reaction. A 100 μL aliquot of protoplast suspension in TMSC per reaction was incubated on ice for 15 minutes. RNP complex, donor template DNA and protoplasts were combined, gently mixed, and incubated on ice for 10 minutes. Subsequently, hand-warm 60% PEG solution (0.6 g/mL PEG 3350, 1 M sorbitol, 10 mM MOPS, pH 6.3) was added, mixed gently, and incubated for 20 min at room temperature. Protoplasts were washed with TMSC solution, resuspended in regeneration medium, and plates as described above.

### Colony propagation and genotyping

Transformant colonies, which appeared at the surface of the regeneration plates, were excised and transferred to 2 mL microcentrifuge tubes containing two 2 mm glass beads. Mycelium was homogenized in sterile water using a TissueLyser II (Qiagen) as described above. The transformant strains were propagated on PDA plates and incubated at 25 °C in the dark.

For genotyping, mycelium was scraped from plates, flash-frozen in liquid nitrogen, and freeze-dried. Lyophilized mycelium was ground using a TissueLyser II (Qiagen). Genomic DNA was extracted using the MasterPure Yeast DNA Purification Kit (BioSearch Technologies) according to the manufacturer’s protocol. Genotyping PCRs were performed using primers listed in Supplemental Table S1.

### Plant growth conditions and infection assays

Cavendish banana (*Musa acuminata* ‘Grand Naine’) and *Musa acuminata* subsp. *burmannicoides* ‘Calcutta 4’ plants, obtained from the International *Musa* Germplasm Transit Center (ITC) were grown under controlled greenhouse conditions (25 °C day/23 °C night, 16 h photoperiod, 80% relative humidity). For infection assays, mycelium from 2-week-old cultures was scraped from plates and homogenized in sterile water using a laboratory blender, filtered through cheesecloth, and adjusted to an OD_600_ of 2.0. Tween 20 was added to a final concentration of 0.1% (v/v). Eight-week-old plants were inoculated by spraying both the adaxial and abaxial side of the leaves until runoff. The inoculum was allowed to dry after which a second spray was applied. Plants were incubated in sealed transparent containers to maintain 100% humidity. Disease progress was assessed at 5, 7, and 9 weeks post inoculation. Disease severity was scored using a 0-5 scale, where 0 corresponded to leaves with no visible symptoms and 5 corresponded to leaves showing complete necrosis.

### Statistical analysis

Statistical analyses were performed using GraphPad Prism. Details of statistical tests are provided in the corresponding figure legends. Data are represented as mean with standard error.

## Results

### Optimization of *P. fijiensis* protoplast generation

In this study, we developed a CRISPR/Cas9-mediated transformation procedure tailored for the fungus *P. fijiensis*. Since the delivery of DNA and RNP complexes into fungal cells is commonly achieved through protoplasts, our initial focus was to establish a robust protocol for high-yield protoplast production.

We first optimized culture conditions to obtain fresh mycelium of *P. fijiensis*, which serves as input biomass for protoplasting. The evenly dispersed hyphal growth observed in potato dextrose broth (PDB) after 40 h was identified as the most favorable condition because it allowed for uniform exposure to cell wall-degrading enzymes, and, therefore, potentially more efficient protoplasting (Figure 1A). In contrast, other culture media resulted in clumped, aggregated hyphal growth, potentially hindering cell wall-degrading enzyme accessibility and therefore reducing protoplasting efficiency (Supplemental Figure S1A).

**Figure 1.**
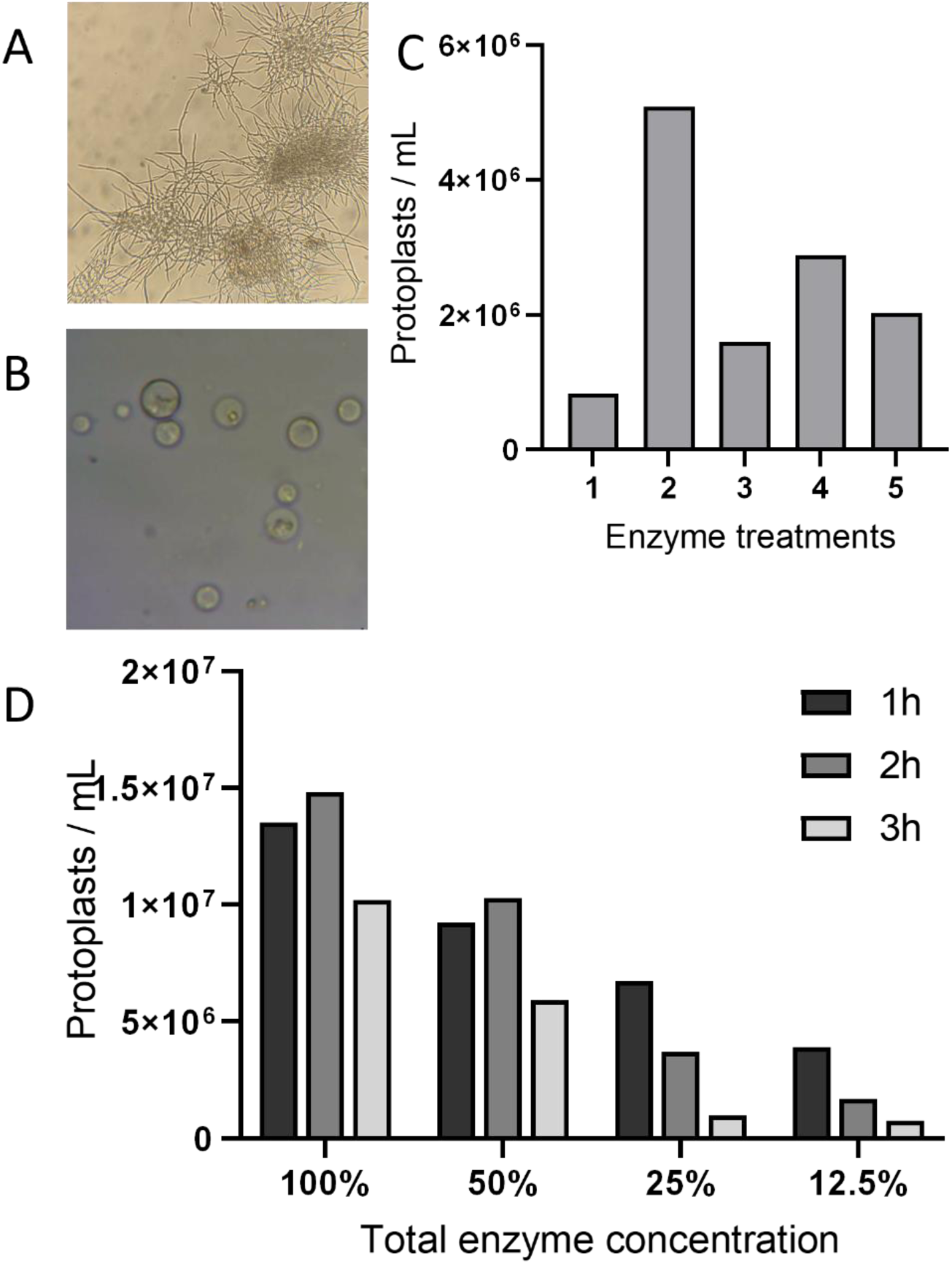
Optimization of protoplast generation in *P. fijiensis*. **A.** Hyphal growth of *P. fijiensis* after 48 hour cultivation in Potato Dextrose Broth (PDB), showing evenly dispersed mycelium suitable for protoplasting. **B.** Representative image of *P. fijiensis* protoplasts following enzymatic digestion. **C.** Protoplast yield obtained after enzymatic treatment of *P. fijiensis* hyphae with different combinations of cell wall-degrading enzymes. Enzyme mixtures were based on protoplasting protocols described for various filamentous fungi as described in Table 1: 1. *B. cinerea* (Leisen et al., 2020), 2. *P. fijiensis* (Arango Isaza et al., 2016), 3. *C. fulvum* (Collemare et al., 2014), 4. *P. fijiensis* (Balint-Kurti et al., 2001), 5. *Fusarium oxysporum* (Dijkstra et al., 2025). **D.** Effect of reduced enzyme concentrations and incubation time on protoplast yield.

Next, we evaluated cell wall-degrading enzymes for their protoplasting efficacy. Five enzyme combinations, based on established protoplasting protocols for other filamentous fungi, were evaluated, including two protocols previously used for *P. fijiensis* (Table 1). Our assessment demonstrated that the enzyme mixture containing Yatalase, Glucanex, β-glucoronidase and Driselase, previously described by Arango Isaza et al. (2016), yielded the highest number of protoplasts (Figure 1B, C).

To improve the efficiency and cost-effectiveness of the protoplasting procedure, we subsequently assessed the individual protoplasting capacity of these enzymes. Although each tested enzyme could generate protoplasts to some extent, their individual contribution to the overall yield was limited (Supplemental Figure S1B). We further investigated the contribution of individual enzymes by systematically omitting them from the total enzyme mixture as described by Arango Isaza et al. (2016) (Supplemental Figure S1C). We concluded that all enzymes contributed in concert to protoplast formation, with β-glucoronidase having the smallest effect.

To further optimize the protoplasting efficiency, we assessed the ratio between enzyme concentration and biomass input. Notably, a 50% reduction in total enzyme concentration maintained a robust protoplast yield suitable for downstream transformation (Figure 1D). Finally, we determined the optimal incubation time for protoplasting to be 2 hours (Figure 1D).

### Selection marker sensitivity and protoplast regeneration into fungal colonies

Following protoplasting optimization, we assessed their ability to regenerate into new fungal colonies and internalize and integrate foreign DNA. Firstly, the sensitivity of protoplasts to the antibiotic selection marker hygromycin B was evaluated. Protoplasts were regenerated in regeneration medium for 18 hours, after which they were covered with an overlay containing a concentration gradient of hygromycin B. Control plates were covered with an overlay without the selection marker. We determined that the minimal effective selection threshold was 75 µg hygromycin B/mL in the overlay (Figure 2A). Plates containing low concentrations of hygromycin B showed a patchy distribution of colonies, likely induced due to local variation in overlay thickness, whereas the controls were completely covered with regenerated colonies.

**Figure 2.**
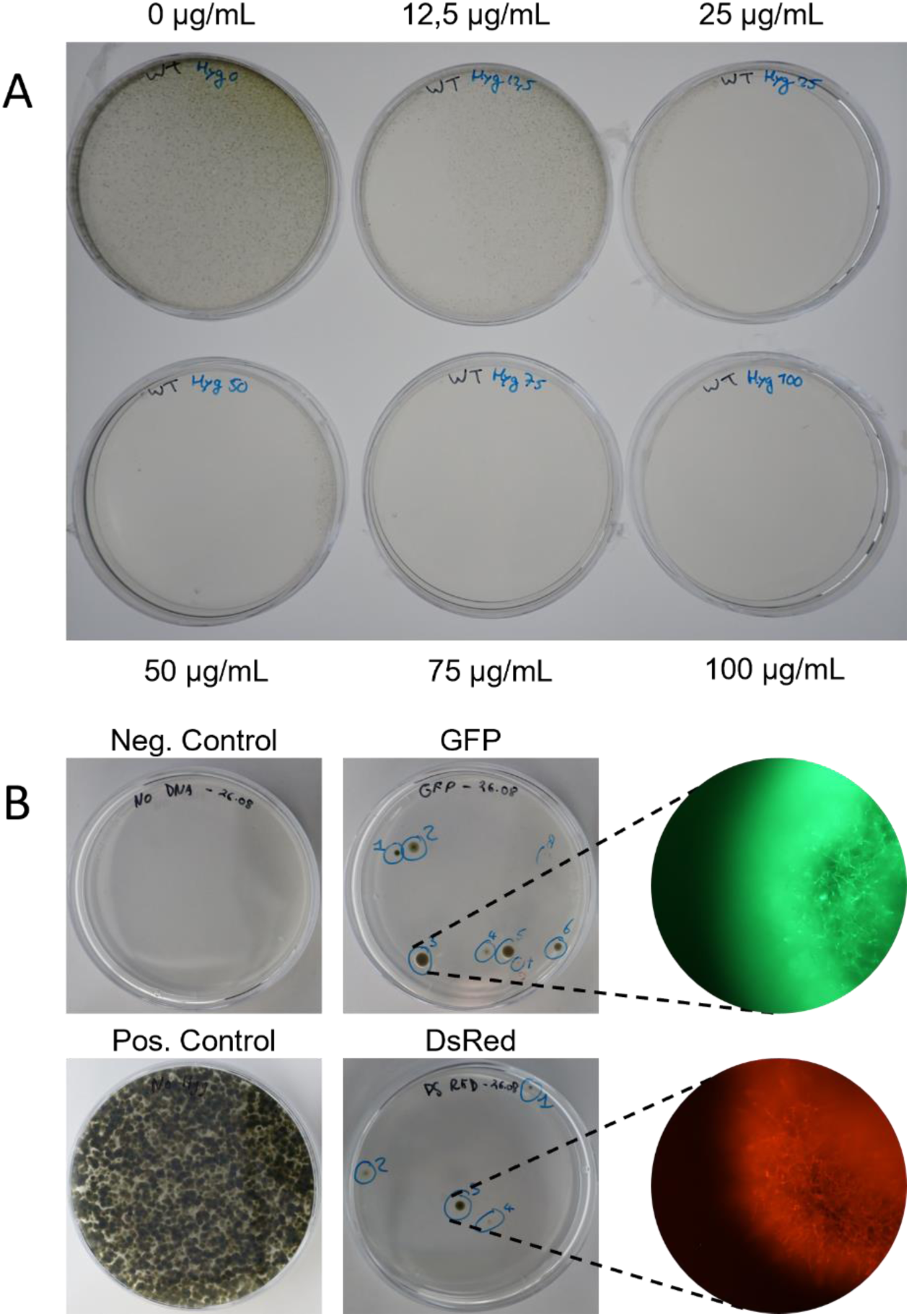
Protoplast selection and regeneration **A.** Colony formation of *P. fijiensis* on plates containing a hygromycin B concentration gradient, 7 days after protoplast regeneration. Indicated concentrations correspond to the antibiotic concentration in the overlay. **B.** Colony formation after PEG-mediated transformation of *P. fijiensis* protoplasts using constructs containing a hygromycin B resistance cassette and either GFP or DsRed as fluorescent reporters, 2 weeks after plating. Representative close-up images of a single fluorescing colony per transformation are shown.

To assess the uptake and incorporation of foreign DNA into the genome of *P. fijiensis*, PEG-mediated transformation was performed. Protoplasts were transformed with constructs lacking homology arms but containing a hygromycin B resistance cassette and a fluorescent reporter gene. Both GFP and DsRed constructs were successfully internalized by the protoplasts and ectopically integrated into the genome. Fifteen and six colonies were recovered following transformations with GFP and DsRed constructs, respectively, and their fluorescence was assessed by microscopy (Figure 2B). Of these, 66% of GFP and 83% of DsRed transformants exhibited fluorescence, indicating successful integration and expression of the respective constructs. The absence of fluorescence in a small subset of transformants may be explained by partial integration events, in which fragments of the construct, such as the reporter gene, are lost or rearranged during transformation. These results demonstrate that PEG-mediated transformation is an effective method for introducing foreign DNA into *P. fijiensis* protoplasts and confirm the suitability of hygromycin B as a selection marker. Clearly, the fluorescently labeled strains are invaluable for detailed microscopic studies on the pathogenesis of *P. fijiensis* in banana foliage.

### CRISPR/Cas9-mediated transformation targeting *PKS10-1* and *Fus3*

To evaluate the efficiency of CRISPR/Cas9-mediated genome editing in *P. fijiensis*, we selected *PKS10-1* and *Fus3* for generating knock-out mutants. Disruption of the polyketide synthase gene *PKS10-1*, which is essential for biosynthesis of the pigment melanin (Churchill, 2011; Noar and Daub, 2016), would enable straightforward visual screening based on colony pigmentation. The mitogen-activated protein kinase (MAPK) gene *Fus3* has a conserved role in plant pathogenic fungal processes such as host penetration, invasive growth, and virulence (Cousin et al., 2006; Onyilo et al., 2018), as well as more fundamental aspects such as growth and developmental differentiation processes (Xu, 2000). Therefore, we hypothesized that disruption of *Fus3* in *P. fijiensis* would lead to reduced virulence during infection.

For each gene, two sgRNAs were designed to target distinct regions within the coding sequence (Figure 3A). The efficacy of each sgRNA in guiding the RNP complex to the target locus and inducing a double-strand break was validated through an *in vitro* cleavage assay. PCR-amplified target fragments were incubated with pre-assembled Cas9-sgRNA RNP complexes, and the cleavage products were analyzed by gel electrophoresis. For *PKS10-1*, both sgRNAs efficiently induced double-strand breaks at the expected positions, yielding fragments of predicted sizes (5650 bp and 1980 bp for sgRNA1, and 3827 bp and 3803 bp for sgRNA2, respectively) (Figure 3B).

**Figure 3.**
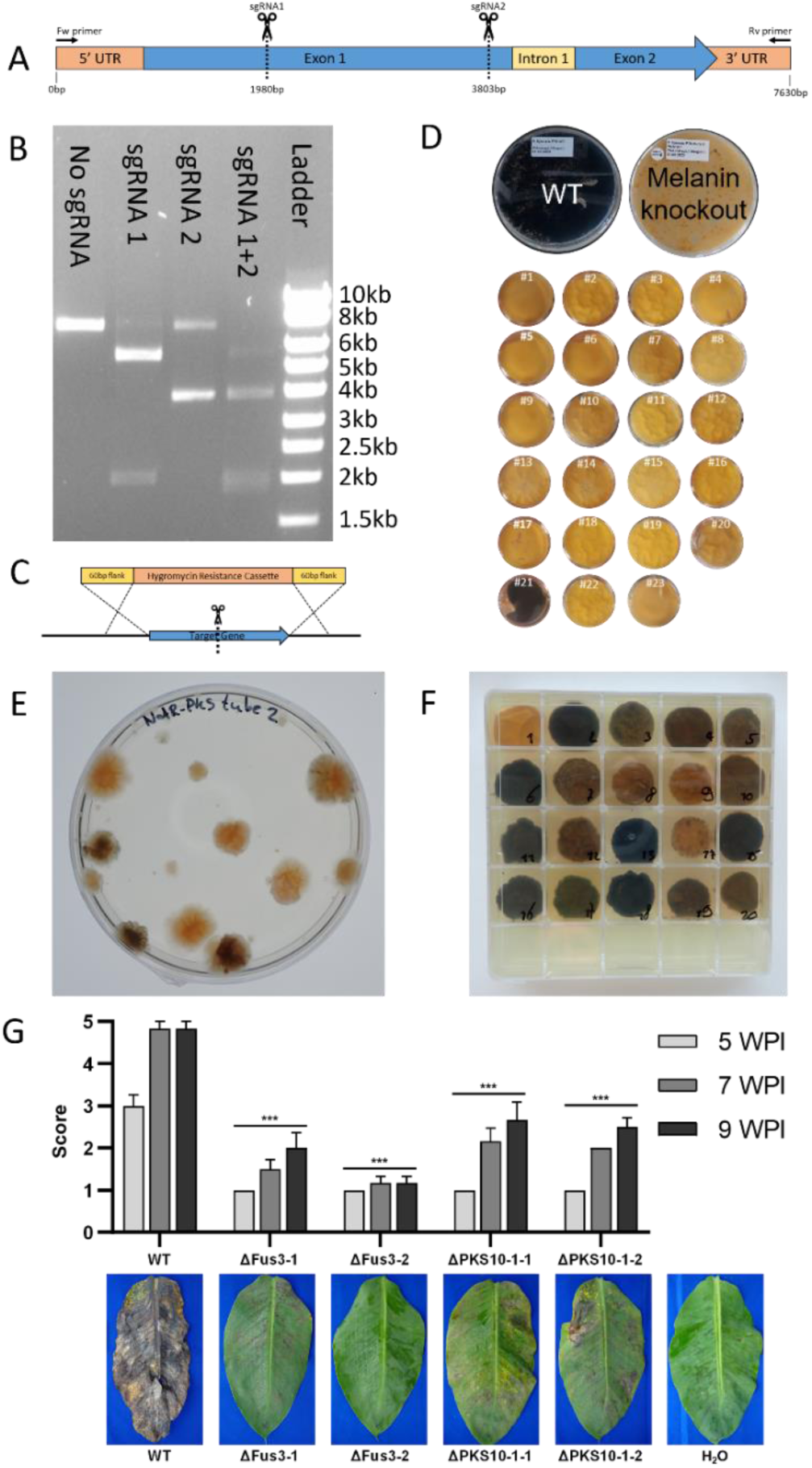
CRISPR/Cas9-mediated transformation of *P. fijiensis*. **A.** Schematic representation of *PKS10-1* target DNA amplicon with the Cas9 cleavage sites (dashed lines) targeted by two sgRNAs. **B.** Gel electrophoresis of DNA fragments from *in vitro* cleavage assay of *PKS10-1* target DNA incubated with Cas9 and the indicated sgRNAs. **C.** Schematic overview of HDR-mediated integration of the donor template following Cas9-induced double-strand break, illustrating integration of a hygromycin B resistance cassette flanked by homology arms. **D.** Colony phenotypes of WT *P. fijiensis* and 23 independent *PKS10-1* transformants **E.** Regenerated colonies following complementation of *PKS10-1* mutants on a selective plate. **F.** Phenotypes of propagated *PKS10-1* complementation lines, showing restoration of pigmentation to varying degrees. **G.** Infection assay of WT *P. fijiensis* and two independent mutant lines for both *Fus3* and *PKS10-1* on banana plants cv. Cavendish. Disease severity was quantified using a disease index at 5, 7, and 9 weeks post-inoculation (WPI). Error bars represent standard error (n=6 plants per strain). Statistical significance between WT and mutant lines was assessed using unpaired t-tests at each time point (*** indicating P≤0.001).

**Figure 4.**
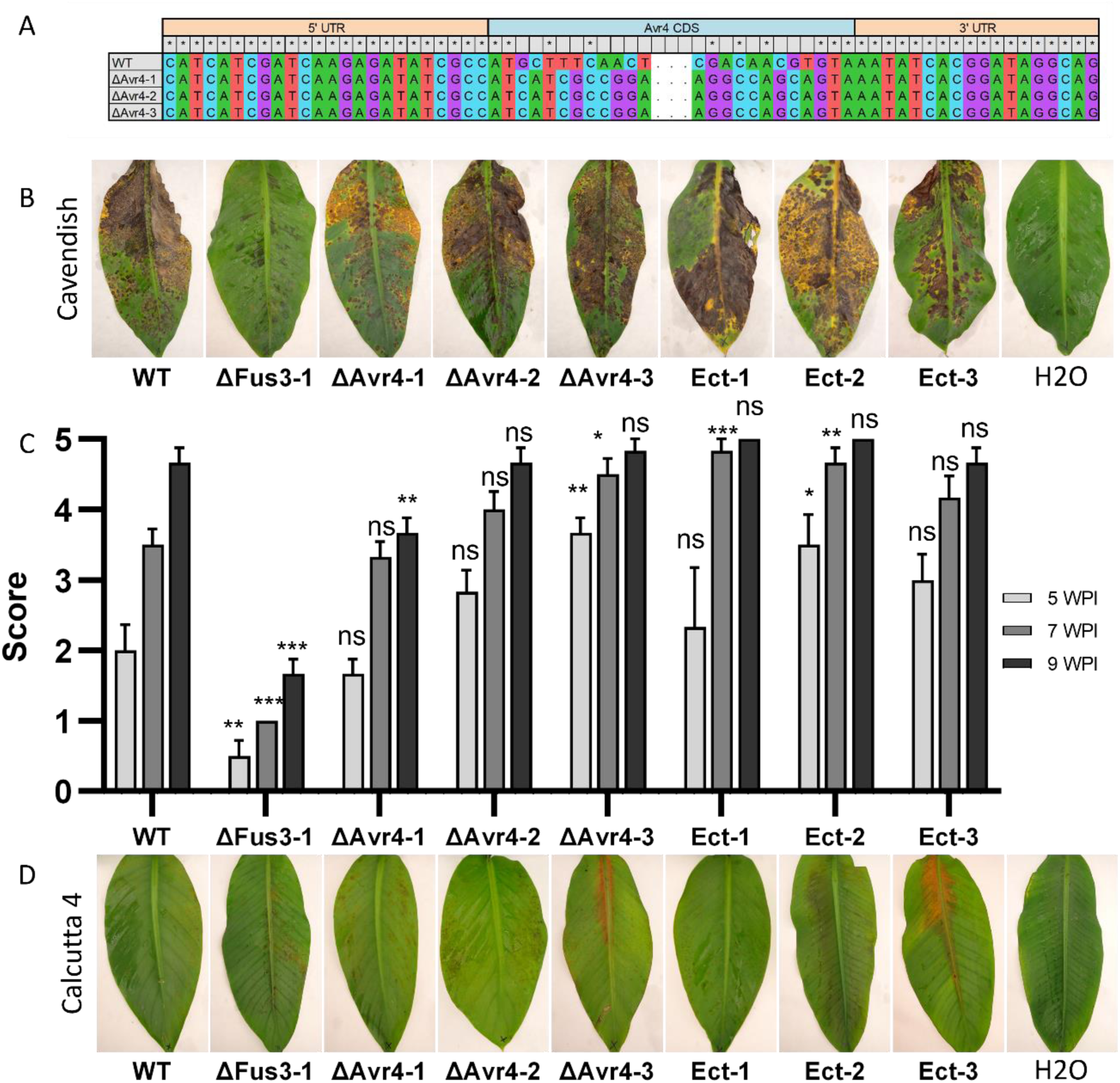
CRISPR/Cas9-mediated knockout of *Avr4* in *P. fijiensis* and its role in virulence and host resistance. **A.** Sequence alignment of the *Avr4* target locus in the wild type and knockout mutants, showing replacement of the coding sequence. **B.** Infection assay of wild type *P. fijiensis*, *Avr4* knockout mutants, and control strains on Cavendish banana plants, showing disease symptom development at 7 weeks post-inoculation (WPI) **C**. Quantification of disease severity on Cavendish plants using a disease index at 5, 7, and 9 WPI. Error bars represent the standard error (n=6 plants per strain). Statistical Differences between wild type and mutant lines were assessed using unpaired t-tests at each timepoint (*** indicating *P*≤0.001) **D.** Infection assay of wild type *P. fijiensis*, *Avr4* knockout mutants, and control strains on the resistant banana cultivar Calcutta 4, showing absence of disease symptoms at 7 WPI.

CRISPR/Cas9-mediated transformations were performed using protoplasts in combination with donor templates containing 60 bp homology arms flanking a hygromycin B resistance cassette (Fig 3C). Transformation yielded dozens of colonies per target gene, which were subsequently propagated. Indeed, disruption of *PKS10-1* resulted in a clear and consistent phenotype that was easily and visually evaluated by the loss of pigmentation. Specifically, 22 out of 23 transformants displayed a yellow, melanin-deficient phenotype, corresponding to a gene-targeting efficiency of 96% (Figure 3D).

PCR-based genotyping revealed that the single pigmented transformant (#21) represented an ectopic integration event, retaining an intact endogenous *PKS10-1* gene (Supplemental Figure S2), whereas all non-pigmented transformants showed disruption of the target gene. Further genotyping analysis indicated that most transformants did not arise from HDR events in which the target locus was fully replaced by the donor template. Instead, they resulted from integration of the donor template at the Cas9-induced double-strand break, likely mediated by non-homologous end joining (NHEJ). Nonetheless, both types of integration resulted in an effective disruption of *PKS10-1*, which was confirmed by complementation experiments. Sensitivity of the protoplasts to a second selection marker, nourseothricin, was first determined, with a minimal effective concentration of 50 µg/mL in the overlay (Supplemental Figure S3). A functional copy of *PKS10-1* was fused to a nourseothricin resistance cassette and introduced into a melanin-deficient strain via ectopic transformation. Transformants displayed a range of pigmentation phenotypes, from yellow to fully melanized (Figure 3E). Upon propagation of the transformants, these differences became more pronounced with colonies exhibiting distinct pigmentation, ranging from non-pigmented (colony 1, 9, 17), to partially pigmented (colony 3, 5, 7) and fully restored melanin production (colony 13, 15, 18) (Figure 3F). This variation in pigmentation can likely be explained by differences in the expression levels of the ectopically integrated *PKS10-1*. The differences in expression may potentially result from positional effects and the use of its native promoter outside of the endogenous secondary metabolite gene cluster, where expression is normally tightly regulated (Brakhage, 2013; Keller, 2019).

For the CRISPR/Cas9-mediated transformation of *Fus3*, the majority of transformants exhibited reduced growth and the expected altered colony morphology. PCR-based genotyping of twelve propagated transformants showed that for seven strains the *Fus3* gene was disrupted, corresponding to a gene-targeting efficiency of 58%. (Supplemental Figure S4). Similar to the *PKS10-1* transformants, most *Fus3* mutants resulted from donor template integration at the double-strand break rather than full replacement of the target locus by HDR, but nonetheless led to functional gene disruption.

To assess the impact of *PKS10-1* and *Fus3* disruption on pathogenicity, infection assays were performed. Two independent knockout lines were selected for both *PKS10-1* and *Fus3*. Disease progression was monitored and scored at 5, 7, and 9 weeks post inoculation. All *PKS10-1* and *Fus3* knockout lines showed significantly attenuated disease symptoms compared to the wild type across all time points (Figure 3G). The reduced virulence of *Fus3* mutants is consistent with previous observations (Onyilo et al., 2018)(Onyilo et al. 2018), whereas the phenotype observed for *PKS10-*1 differs from earlier reports on the role of melanin in *P. fijiensis* virulence (Churchill, 2011).

### Assessing the role of *Avr4* in virulence and host resistance

*Avr4* is one of only two effectors thus far characterized in *P. fijiensis* (Stergiopoulos et al., 2010). To elucidate its role in fungal virulence, we generated *Avr4* knockout mutants using the CRISPR/Cas9-mediated approach. Similar to the strategy used for generating mutants of *PKS10-1* and *Fus3*, a donor template was constructed containing a hygromycin B resistance cassette flanked by 60 bp homology arms corresponding to the upstream and downstream regions of the *Avr4* coding sequence. Following transformation, 31 colonies were obtained, of which 20 were propagated for further analysis. None of the transformant colonies displayed morphological differences compared to the wild-type. PCR-based genotyping revealed successful donor template integration at the correct genomic locus for 11 out of the 20 screened colonies (Supplemental Figure S5), corresponding to a 55% gene-targeting efficiency. Analogous to *PKS10-1* and *Fus3*, the majority of the *Avr4* mutants arose from NHEJ-mediated donor template integration rather than HDR. Although both integration events resulted in disruption of *Avr4*, we specifically selected HDR-mediated full replacement mutants for downstream analysis. These mutants were confirmed by sequence analyses (Figure 3A), excluding the possibility that truncated Avr*4* fragments, while non-functional, could still be expressed and recognized by host immune receptors during plant infection.

The disease assays included the wild-type strain, three independent full replacement *Avr4* mutants, three ectopic integration controls, along with a *Fus3* knockout mutant, which showed reduced virulence. Disease progression was monitored and scored at 5, 7, and 9 weeks post inoculation. Neither the *Avr4* knockout mutants nor the controls exhibited any reduction in virulence compared to the wild-type at any time point, indicating that *Avr4* is not required for full virulence on Cavendish (Figure 3B, C).

Arango Isaza et al. (2016) infiltrated heterologously produced Avr4 protein from *P. fijiensis* into the resistant banana cultivar Calcutta 4, which triggered a cell death response, suggesting a potential role for *Avr4* in the resistance of Calcutta 4 by host recognition. To test this hypothesis, we also inoculated Calcutta 4 with the *Avr4* knockout mutants. Despite the absence of *Avr4*, no disease symptoms were observed, and plants remained fully resistant against the *Avr4* replacement mutants (Figure 3D). Furthermore, hypersensitive response-like symptoms were detected upon inoculation with both mutant and control strains. Together, these results indicate that the resistance of Calcutta 4 to *P. fijiensis* is not driven by the recognition of Avr4.

## Discussion

The limited availability of efficient genetic transformation methods for *P. fijiensis* has long constrained functional genetic studies aimed at elucidating the molecular basis of BLSD. In this study, we developed an efficient CRISPR/Cas9-mediated transformation platform for *P. fijiensis* by systematically optimizing protoplast generation, transformation conditions, and targeted gene disruption. The resulting protocol consistently yielded high numbers of transformants across three tested genes and achieved targeted gene disruption efficiencies of up to 96%, providing a powerful tool for reverse genetic studies in this important banana pathogen.

The ability to efficiently generate targeted gene knockouts in *P. fijiensis* provides new opportunities for systematic functional characterization of candidate effectors and other pathogenicity-associated genes. Similar reverse genetic approaches in other fungal pathosystems, including *Zymoseptoria tritici* and *Parastagonospora nodorum*, have greatly accelerated the identification of virulence and avirulence factors and facilitated the subsequent discovery of corresponding host resistance and susceptibility genes (Liu et al., 2009; Zhong et al., 2017; Meile et al., 2018; Kariyawasam et al., 2022; Richards et al., 2022). The availability of robust reverse genetics resources has contributed to the establishment of these species as model systems for studying fungal pathogenicity and host-pathogen interactions (Oliver et al., 2012; Kariyawasam et al., 2023; Meile et al., 2025). Establishing similar reverse genetic strategies to *P. fijiensis* will enable the dissection of molecular mechanisms underlying pathogenicity and host specificity. This knowledge will contribute to the identification of resistance genes and support the development of more durable disease-control strategies for banana cultivation (Drenth and Kema, 2024).

Successful transformation of many filamentous fungi relies on the generation of large numbers of viable protoplasts. By optimizing culture conditions, enzyme composition, enzyme concentration, and incubation time, we established a reproducible and efficient protocol for protoplast generation in *P. fijiensis*. The enzyme mixture adapted from Arango Isaza et al. (2016) proved highly effective in generating viable protoplasts suitable for transformation. However, this enzyme combination consists of several specialized enzymes that, in some cases, may not always be readily available. Recent studies in other filamentous ascomycetes, including *A*. *nidulans*, *Colletotrichum lindemuthianum*, and *B*. *cinerea* (Oakley et al., 2012; Coca Ruiz et al., 2024; Nabi et al., 2025), have reported successful protoplast generation using alternative enzyme blends such as Vinotaste Pro, a commercially available enzyme mixture originally developed for winemaking. These enzyme preparations may represent accessible and cost-effective alternatives, although their performance in *P. fijiensis* remains to be evaluated. Future studies may assess whether such enzyme formulations can further facilitate large-scale protoplast generation and transformation, particularly for high-throughput genetic analyses.

In this study, we used hygromycin B and nourseothricin as selection markers during transformation. Expanding the repertoire of selection markers would further increase the utility of the transformation system. The availability of multiple independent selection markers would enable sequential rounds of gene editing, allowing the generation of strains carrying multiple targeted mutations. However, multiplex CRISPR/Cas9 approaches, in which several sgRNAs and donor templates are delivered simultaneously, may enable the disruption of multiple genes in a single transformation event, as has been demonstrated in model fungi such as *Saccharomyces*, *Aspergillus*, and *Trichoderma* (Ryan et al., 2014; Liu et al., 2015; Nødvig et al., 2018). Sensitivity to selection markers may vary among *P. fijiensis* isolates. This is particularly relevant as field populations have developed reduced sensitivity to several fungicides used in commercial banana production, including compounds belonging to the chemical class of sterol demethylation inhibitors (DMIs), strobilurins and benzimidazols (Chong et al., 2021; Manzo-Sánchez et al., 2024; Maridueña-Zavala et al., 2024). Such reduced sensitivity could potentially also influence their sensitivity to antibiotic selection and should therefore be considered when applying selection markers in transformation experiments involving genetically diverse field isolates.

The melanin biosynthesis gene *PKS10-1* was selected as a benchmark target to evaluate the efficiency of the developed CRISPR/Cas9-mediated transformation system. This gene encodes a polyketide synthase that is essential for melanin biosynthesis, and disruption of this gene resulted in an easily distinguishable unpigmented yellow-colored phenotype, consistent with previous reports (Churchill, 2011; Noar and Daub, 2016). Melanin has been implicated in multiple aspects of fungal biology and pathogenicity. In several pathogenic fungi, melanin plays a direct role in host infection. For example, in the model fungus *M*. *oryzae*, melanin is an essential component of the appressorium, where it facilitates the accumulation of glycerol and the generation of the high turgor pressure required for penetration of the host plant surface (Howard and Ferrari, 1989). In addition, melanin has been associated with protection against ultraviolet radiation, increased cell wall integrity, and protection against reactive oxygen species (ROS) during plant defense responses (Wang and Casadevall, 1994; Nosanchuk and Casadevall, 2006; Noar et al., 2022; Jia et al., 2025). In *P. fijiensis*, melanin has also been proposed to function as a light-activated phytotoxin capable of generating ROS that contribute to host tissue damage during infection (Beltran-Garcia et al., 2014). In contrast to a study reporting that disruption of melanin biosynthesis did not affect pathogenicity in *P. fijiensis* (Churchill, 2011), our *PKS10-1* knockout mutants exhibited a clear reduction in virulence compared to the wild type. The reasons for this discrepancy remain unclear but may reflect differences in the genetic background of the strains used, inoculation procedures or environmental conditions during disease development. It is also possible that melanin contributes to pathogenicity in a context-dependent manner, for example under conditions that promote oxidative stress or increased light exposure. Alternatively, the reduced virulence observed in our mutants may not be solely attributable to the absence of melanin itself. Disruption of the melanin biosynthetic pathway can alter the production of secondary metabolites derived from pathway intermediates through so-called melanin shunt metabolism (Stierle et al., 1991; Hoss et al., 2000; Churchill, 2011; Noar et al., 2022). Several of these metabolites, including the phytotoxins juglone and fijiensin, have been extensively characterized in *P. fijiensis* (Upadhyay et al., 1990; Stierle et al., 1991). Disruption or chemical inhibition of melanin biosynthesis alters the accumulation of melanin shunt metabolites (Hoss et al., 2000; Noar et al., 2022), suggesting that the reduced pathogenicity of the *PKS10-1* disruption mutants may not only result from the loss of melanin itself, but could also reflect altered production of phytotoxic secondary metabolites. Further studies will therefore be required to disentangle the relative contributions of melanin and melanin-derived secondary metabolites to disease development in *P. fijiensis*.

An important observation throughout this study was the high frequency of donor template integration events mediated by non-homologous end joining (NHEJ). Although these events generally resulted in effective gene disruption and were therefore sufficient for knockout generation, certain applications require precise homology-directed repair (HDR). For example, complete gene replacement is preferable when studying effector genes, as residual gene fragments may still be transcribed or translated and subsequently be recognized by plant receptor proteins. Likewise, precise HDR-mediated integration is desirable for gene tagging, promoter replacement, or targeted overexpression studies. Several strategies have been reported to increase HDR frequencies in filamentous fungi, including extending homology arm lengths and suppressing components of the NHEJ pathway. However, the effectiveness of these approaches appears to depend strongly on the involved species. Whereas longer homology arms substantially improved HDR frequencies in some fungi, including *Fusarium* spp. (Shinkado et al., 2022), similar strategies yielded little improvement in others, such as *B. cinerea* (Leisen et al., 2020).

In addition to targeting two reference genes, we also targeted the effector *Avr4* to enable functional characterization and provide insight into the biological role of this effector in the interaction with susceptible and resistant banana accessions. *Avr4* was originally identified and characterized in the tomato pathogen *Cladosporium fulvum*, where it functions as both a virulence factor and an avirulence determinant (Joosten et al., 1994). In susceptible hosts, Avr4 binds chitin in the fungal cell wall and protects the fungus against degradation by plant chitinases (Van den Burg et al., 2006). In resistant tomato cultivars carrying the Cf-4 immune receptor, however, Avr4 is recognized, triggering a defense response that prevents disease development (Thomas et al., 1997). The Avr4 homolog of *P. fijiensis*, PfAvr4, shares 42% amino acid similarity with *C. fulvum* Avr4, including conservation of all cysteine residues that are critical for protein structure (Stergiopoulos et al., 2010). Furthermore, previous work demonstrated that PfAvr4 is recognized by the tomato Cf-4 receptor (Stergiopoulos et al., 2010) and that infiltration of crude filtrate from *Pichia pastoris* expressing recombinant PfAvr4 protein triggered a hypersensitive-like response in the resistant banana accession Calcutta 4 (Arango Isaza et al., 2016). These observations suggested that recognition of Avr4 might contribute to resistance against *P. fijiensis*. However, our results do not support this hypothesis. Deletion of *Avr4* had no measurable effect on fungal virulence on Cavendish banana, indicating that the effector is not required for successful infection under the conditions tested. More importantly, *Avr4* knockout mutants remained completely unable to infect Calcutta 4 and still elicited hypersensitive response-like symptoms upon inoculation. These findings demonstrate that resistance of Calcutta 4 cannot be explained solely by recognition of Avr4. Instead, resistance likely depends on the recognition of one or more additional fungal effectors, either independently or in combination with Avr4. Given the large predicted effector repertoire encoded by the *P. fijiensis* genome (Arango Isaza et al., 2016; Carreón-Anguiano et al., 2024), it is plausible that Calcutta 4 recognizes multiple effectors, resulting in a broad-spectrum and robust resistance response against diverse *P. fijiensis* isolates.

In conclusion, we established a highly efficient CRISPR/Cas9-mediated transformation system for *P. fijiensis* and demonstrated its utility for targeted gene disruption and functional genomics. The platform overcomes a major experimental bottleneck in this notoriously difficult pathogen and provides an essential tool for future studies aimed at unraveling the molecular basis of BLSD and identifying novel sources of resistance for sustainable banana production.

## Acknowledgements

We thank Jelmer Dijkstra and Carolina Aguilera-Galvez for helpful discussions during the development of the project. We are also grateful to Giuliana Nakasato-Tagami for technical assistance. We thank Anouk van Westerhoven and Xiaoqian Shi-Kunne for bioinformatical support. Finally, we would like to thank Matthias Hahn for sharing transformation protocols and plasmids.

