## Supplemental Tables and Figures for "CRISPR/Cas9-mediated transformation enables functional characterization of the effector *Avr4* in the banana pathogen *Pseudocercospora fijiensis*"

### Supplemental Table S1

**Supplemental Table S1.** Oligonucleotides used in this study.

| No. | Name | 5' to 3' sequence |
|---|---|---|
| 1 | sgRNA_constant | AAAAGCACCGACTCGGTGCCACTTTTTCAAGTTGATAACGGACTAGCCTTATTTTAACTTGCTATTTCTAGCTCTAAAAC |
| 2 | Pfij_PKS10-1_specific_sgRNA1 | AAGCTAATACGACTCACTATAGGCAGTATCGACAGAGTACGAGTTTTAGAGCTAGAAATAGCAAG |
| 3 | Pfij_PKS10-1_specific_sgRNA2 | AAGCTAATACGACTCACTATAGGAATTGACAGTCTCACGAGCGGTTTTAGAGCTAGAAATAGCAAG |
| 4 | Pfij_Fus3_specific_sgRNA1 | AAGCTAATACGACTCACTATAGGAGGACATTGGCGGAATGCAGTTTTAGAGCTAGAAATAGCAAG |
| 5 | Pfij_Fus3_specific_sgRNA1 | AAGCTAATACGACTCACTATAGGTACGTTCAAGGAATACACCAGTTTTAGAGCTAGAAATAGCAAG |
| 6 | Pfij_Avr4_specific_sgRNA1 | AAGCTAATACGACTCACTATAGGATCTGCGGACGGGCATACGAGTTTTAGAGCTAGAAATAGCAAG |
| 7 | Pfij_Avr4_specific_sgRNA2 | AAGCTAATACGACTCACTATAGGATACATTCAATGCTCGCCAGGTTTTAGAGCTAGAAATAGCAAG |
| 8 | Pfij_PKS10-1_target_fw | CTGGTCTATCAATGGAACGCG |
| 9 | Pfij_PKS10-1_target_rv | GCCAGCTGTTGATGTATGAAGC |
| 10 | Pfij_Fus3_target_fw | AACGTGTCGCACTAGAGTAGG |
| 11 | Pfij_Fus3_target_rv | CAGAACGACTGGAATTGAGCG |
| 12 | Pfij_Avr4_Target_Fw | TTCCTTCCCTCCCTTCTAGTT |
| 13 | Pfij_Avr4_Target_Rv | TTGCCTCCTTCTTAACCCTAATC |
| 14 | Pfij_PKS10-1_fw_donor_template | ATGTCGAACATCTTACTCTTCGGCGATCAGACCGCCGAGCAATACCCTCTCCTGAGAAAGTGCTGGCCTTTTGCTCACATGCATG |
| 15 | Pfij_PKS10-1_rv_donor_template | GTGGAGTGTAACCTGGTCGGATGGTACCACTCACTTGCACCGTTGATGCTGACTCCTGCTATCGCCGGAAAGGACCCGCAAATG |
| 16 | Pfij_Fus3_fw_donor_template | CGGCTCGCGCAAGATTAGCTTCAACGTATCTGAGCAGTACGACATCCAAGATGTGGTGGGTGCTGGCCTTTTGCTCACATGCATG |
| 17 | Pfij_Fus3_rv_donor_template | TTACCGCATGATCTCCTCATAAATCAGACCTATTCAATCAGCGTCAGTAGAGTTGGAAATATCGCCGGAAAGGACCCGCAAATG |
| 18 | Pfij_Avr4_Fw_donor_template | CTGCGCGCATGGTCAGCATGAATCTCGACATCAAGAAGAAAGCGATCGCCAATCACGTACTGCTGGCCTTTTGCTCACATGCATG |
| 19 | Pfij_Avr4_Rv_donor_template | ACCGATCTCCTAGCCTTCGTGAGCCCGAAAACCCATCATCGATCAAGAGATATCGCCATCATCGCCGGAAAGGACCCGCAAATG |
| 20 | pCT74_Hyg+GFP/dsRed_Fw | GCGCGTAATACGACTCACTAT |
| 21 | pCT74_Hyg+GFP/dsRed_Rv | GCGCAATTAACCCTCACTAAAG |
| 22 | Pfij_Avr4_Fw_check_transformants | GGAATGGAGATGGGTTGGTAAG |
| 23 | Pfij_Avr4_Rv_check transformants | GGCTCTATAGCTCCGCTAGTAT |
| 26 | Avr4_presence_check_Fw | ACGTTGTCGCATCCTGTTT |
| 27 | Avr4_presence_check_Rv | TAACACTGCTCGCCACATTC |

| | | |
|---|---|---|
| 28 | Trans_5end_integr_check_Fw | GCGATGAGAGGTACGCATTTA |
| 29 | Trans_5end_integr_check_Rv | GTATGAGTCACAGCACCGATAC |
| 30 | Trans_3end_integr_check_Fw | GAGAGCCTTCTAGCTTCGAATAC |
| 31 | Trans_3end_integr_check_Rv | CGGGTTTACCTCTTCCAGATAC |
| 32 | Pfij_PKS10-1_Fw | GGGTCATACGCGATTGTTCT |
| 33 | Pfij_PKS10-1_Nat_Rv | ATGTGAGCAAAAGGCCAGCACGCAGTCCTATGTGGATTAAGG |
| 34 | NatR_PKS10-1_Fw | CCTTAATCCACATAGGACTGCGTGCTGGCCTTTTGCTCACATGCATG |
| 35 | NatR_Rv | ATCGCCGGAAAGGACCCGCAAATG |

### Supplemental Figure S1

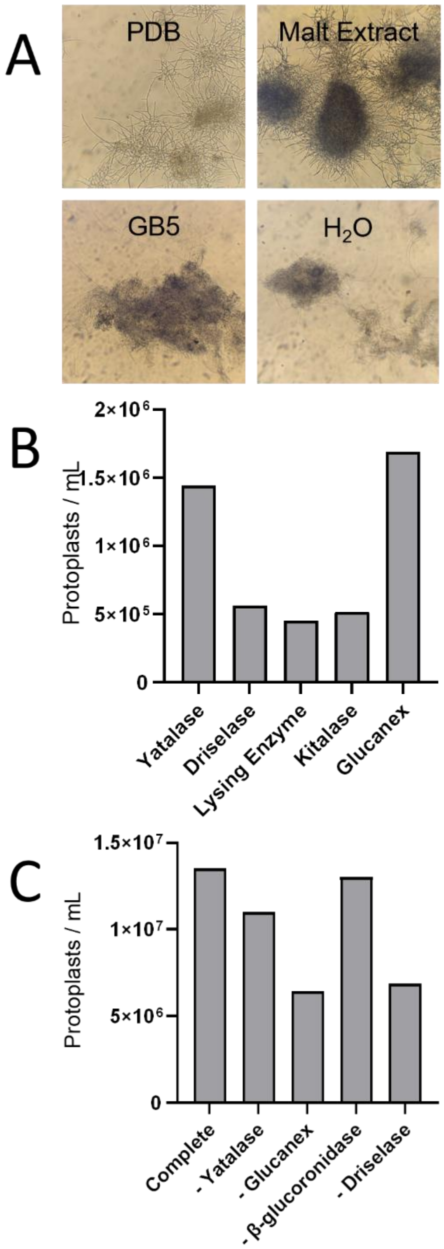


**Supplemental Figure S1.** Optimization of protoplasting generation in *P. fijiensis.* **A.** Hyphal morphology of *P. fijiensis* following growth in different liquid media, showing variation in hyphal dispersion and aggregation. **B.** Protoplast yield obtained using individual cell wall-degrading enzymes, each applied at a final concentration of 10 mg/mL. **C.** Protoplast yield following omission of individual enzymes from the complete enzyme mixture, demonstrating the relative contribution of each enzyme to overall protoplast formation.

### Supplemental Figure S2

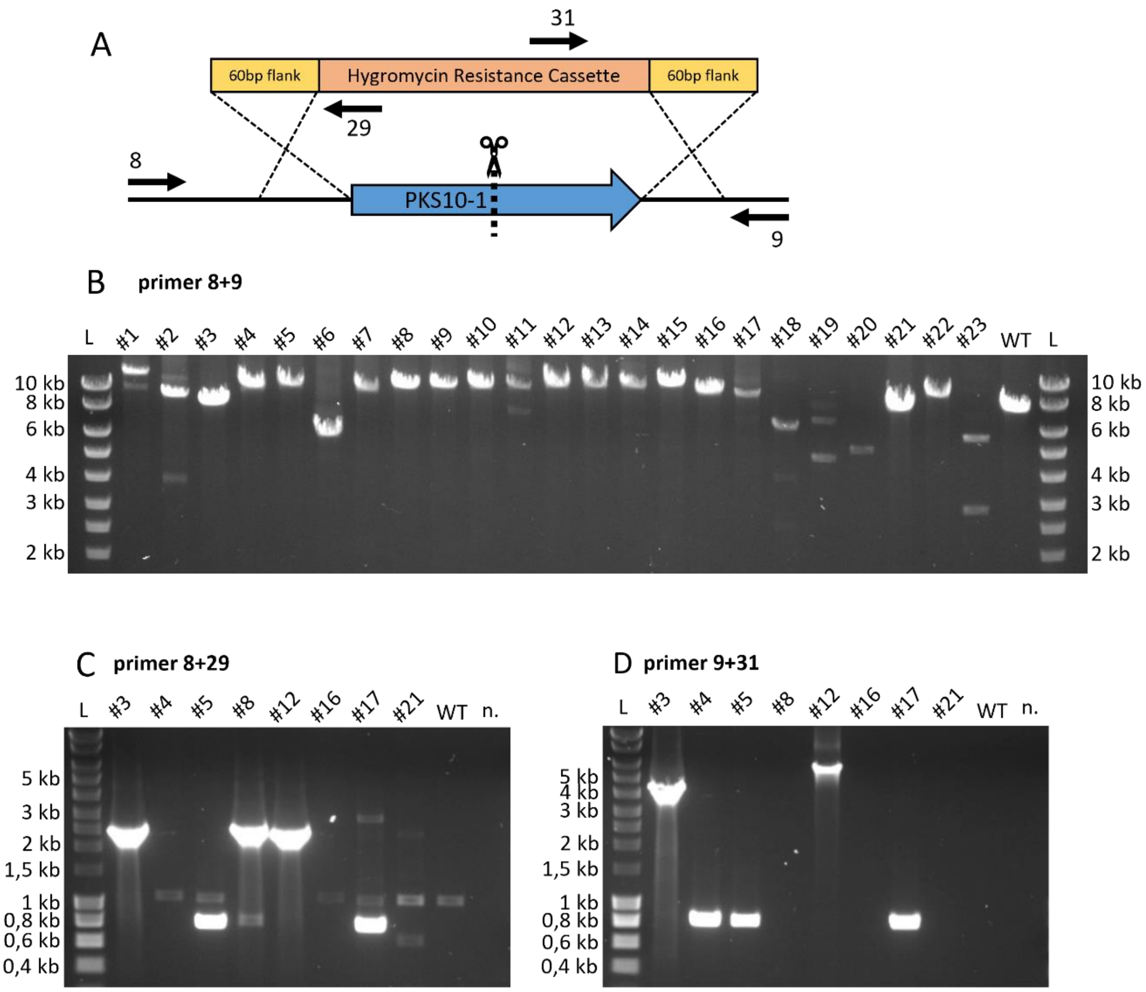


**Supplemental Figure S2.** Molecular genotyping of *PKS10-1* transformants. **A.** Schematic representation of the *PKS10-1* locus and donor template used for transformation. Primer binding sites used for PCR-based genotyping are indicated. **B-D.** Agarose gel electrophoresis of genotyping PCRs using primer pairs indicated in each panel. Transformant colonies are numbered consistently across panels, with WT indicating wild type genomic DNA and n. being a no template control. Primer pairs correspond to those listed in Supplemental Table S1.

### Supplemental Figure S3

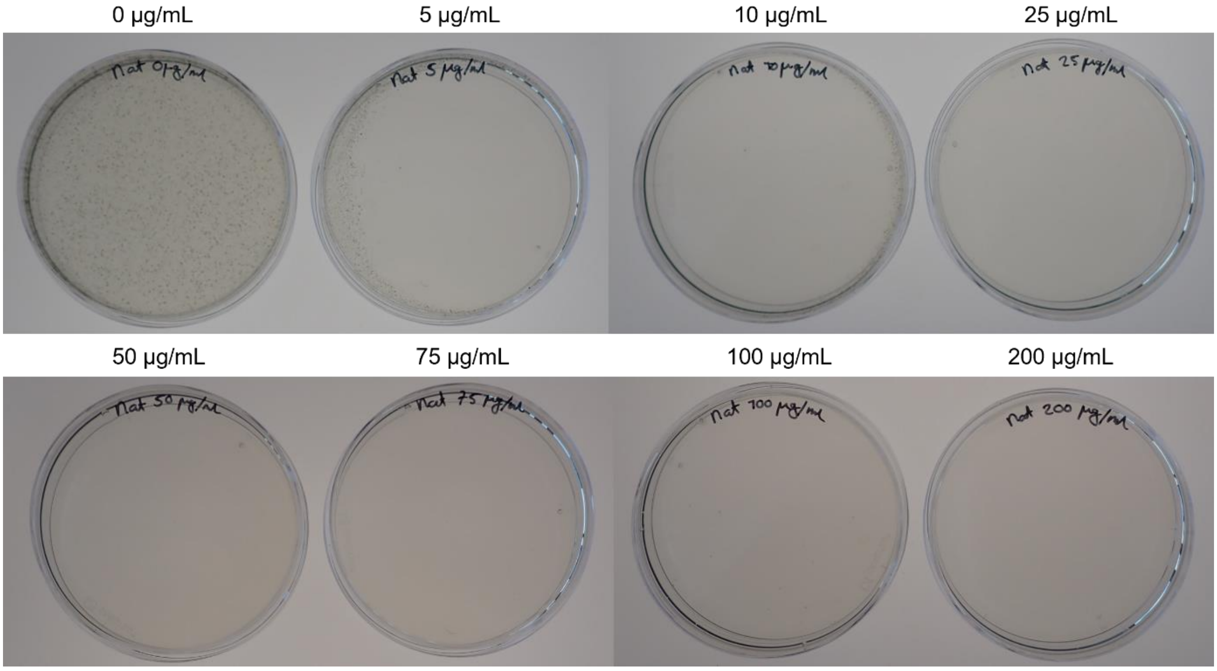


**Supplemental Figure S3.** Colony formation of *P. fijiensis* on plates containing a Nourseothricin concentration gradient, 7 days after protoplast regeneration. Indicated concentrations correspond to the antibiotic concentration in the overlay.

### Supplemental Figure S4

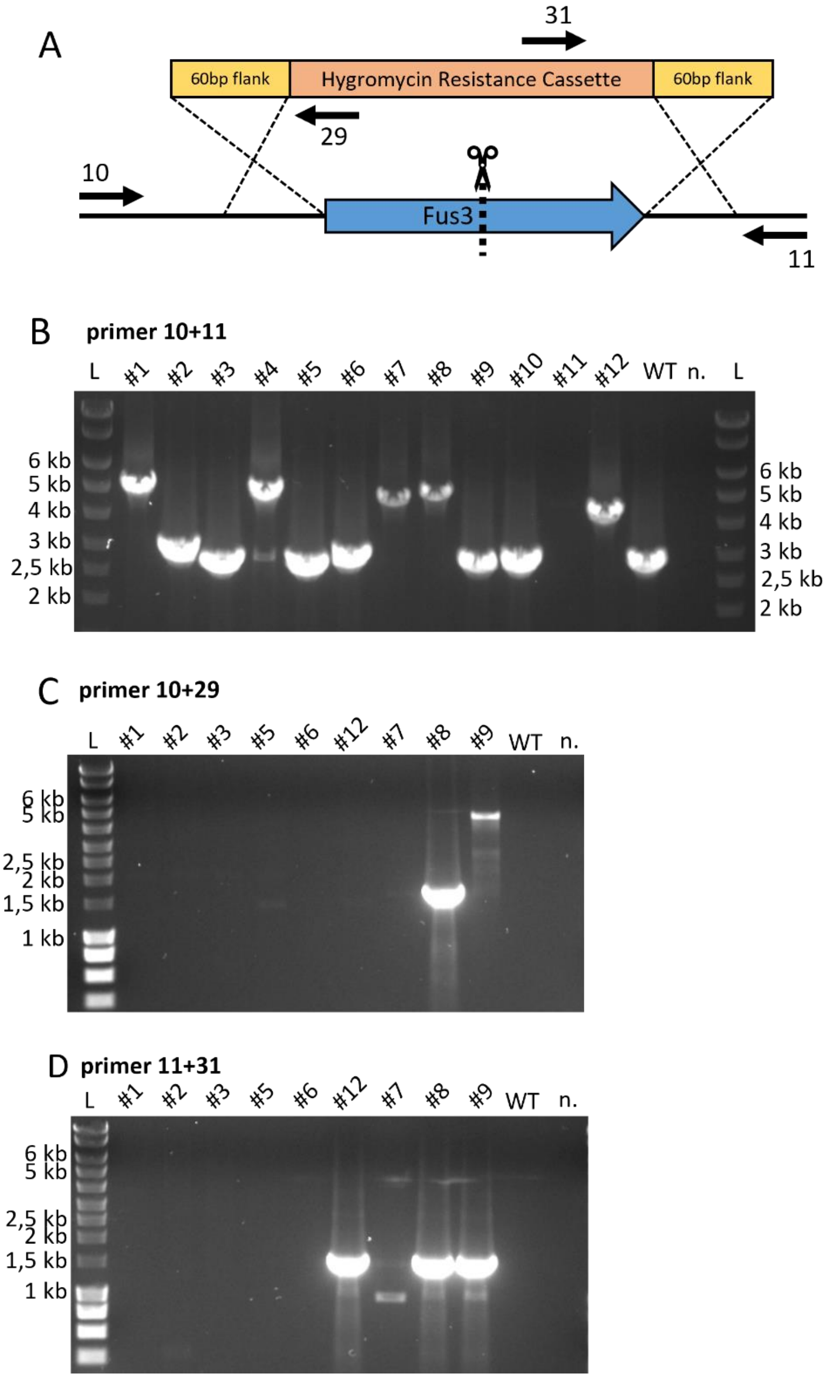


**Supplemental Figure S4.** Molecular genotyping of *Fus3* transformants. **A.** Schematic representation of the *Fus3* locus and donor template used for transformation. Primer binding sites used for PCR-based genotyping are indicated. **B-D.** Agarose gel electrophoresis of genotyping PCRs using primer pairs indicated in each panel. Transformant colonies are numbered consistently across panels, with WT indicating wild type genomic DNA and n. being a no template control. Primer pairs correspond to those listed in Supplemental Table S1.

### Supplemental Figure S5

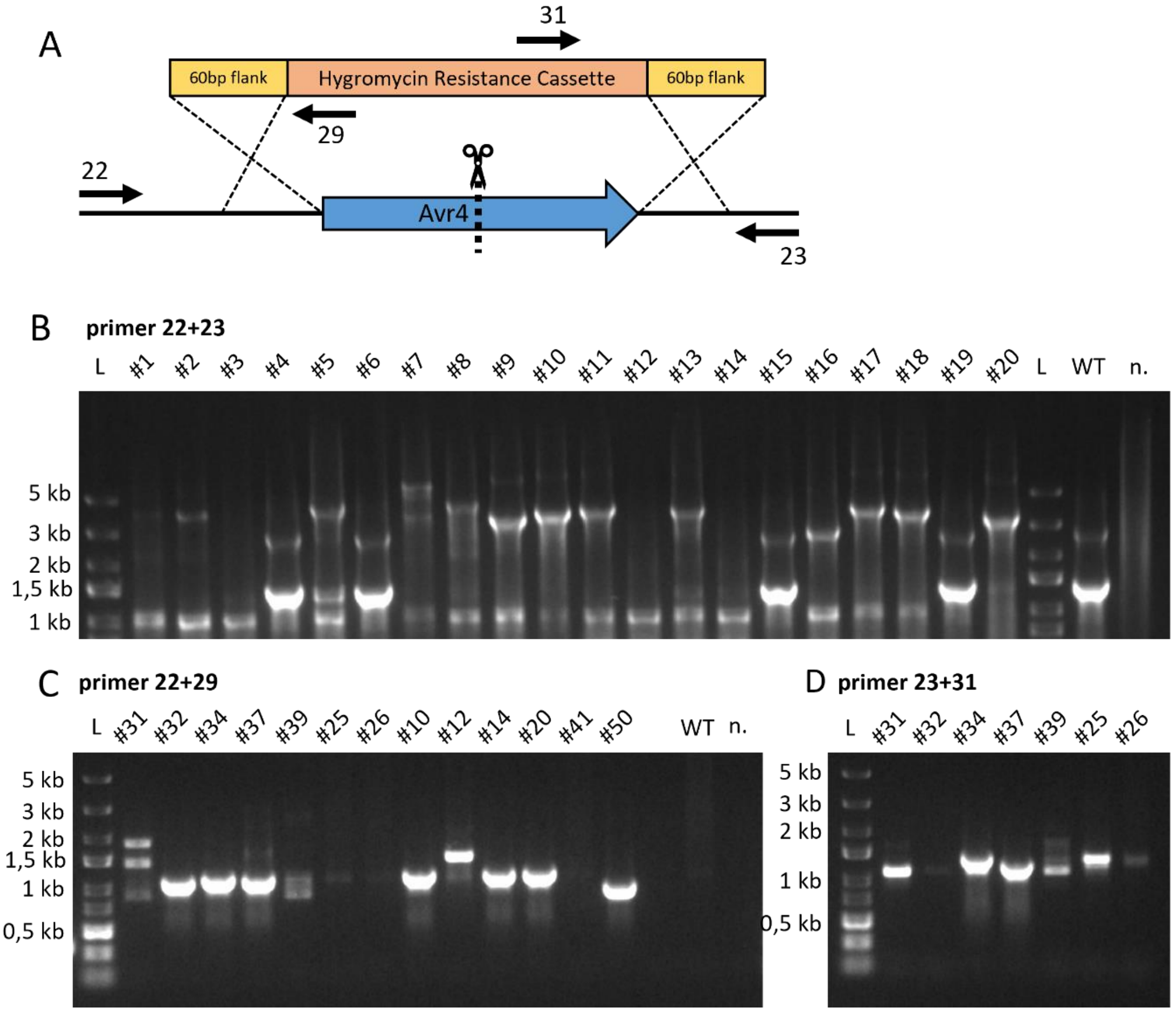


**Supplemental Figure S5.** Molecular genotyping of *Avr4* transformants. **A.** Schematic representation of the *Avr4* locus and donor template used for transformation. Primer binding sites used for PCR-based genotyping are indicated. **B-D.** Agarose gel electrophoresis of genotyping PCRs using primer pairs indicated in each panel. Transformant colonies are numbered consistently across panels, with WT indicating wild type genomic DNA and n. being a no template control. Primer pairs correspond to those listed in Supplemental Table S1.
